# β-Hydroxybutyrate enhances brain metabolism in normoglycemia and hyperglycemia, providing cerebroprotection in a mouse stroke model

**DOI:** 10.1101/2025.02.19.639087

**Authors:** Deborah M Holstein, Afaf Saliba, Damian Lozano, Jiwan Kim, Kumar Sharma, James D Lechleiter

## Abstract

Hyperglycemia in poorly controlled diabetes is widely recognized as detrimental to organ dysfunction. However, the acute effects of hyperglycemia on brain metabolism and function are not fully understood. The potential protective benefit of ketone bodies on mitochondrial function in the brain has also not been well characterized. Here, we evaluated the acute effects of hyperglycemia and β-hydroxybutyrate (BHB) on brain metabolism by employing a novel approach leveraging adenosine triphosphate (ATP)-dependence of bioluminescence originating from luciferin-luciferase activity. Oxygen consumption rate was measured in *ex vivo* live brain punches to further evaluate mitochondrial function. Additionally, we investigated the functional relevance of BHB using an *in vivo* photothrombotic stroke model to assess its cerebroprotective effects. Our data demonstrate that brain metabolism in mice is affected by acute exposure to high glucose, at a level similar to consuming food or a beverage with high sucrose. This short-term effect of glucose exposure was reduced by co-administration with the ketone body BHB. Moreover, BHB significantly reduced infarct size in the brain stroke model, providing evidence for its functional protective role in the brain. These findings suggest that BHB may effectively mitigate the adverse effects of metabolic stress and ischemic events on brain metabolism and function.

## Introduction

The ketogenic diet has been established as a powerful treatment for epilepsy ^1^, and obesity ^2^ and cardiovascular diseases ^3^. Ketones are generated during starvation or states of reduced insulin action and may have beneficial attributes for cells. Ketones have been considered a “super-fuel” for the heart in a diabetic setting ^4, 5^, but it remains unclear what the specific role of low carbohydrate/ketogenic diets is in other energetically demanding organs, such as the brain. Emerging evidence suggests that the macronutrient composition of diets, including low-carbohydrate or ketogenic diets, may influence brain function ^6, 7^. For instance, a clinical trial demonstrated that long-term adherence to high-carbohydrate diets increased regional cerebral blood flow in brain regions associated with hunger and reward, such as the nucleus accumbens and hypothalamus, potentially contributing to weight regain ^8^. In a recent study, we showed that a long-term ketogenic diet increased pro-inflammatory biomarkers in patients and increased brain senescence in mice’s brains, among other organs ^9^. These studies underscore the importance of understanding how macronutrient composition impacts brain energy balance and function, particularly under conditions of metabolic stress.

Glucose is an obligate fuel for neuronal cells, however glucose may be toxic at concentrations above 10 mM ^10^. Several studies demonstrated that high glucose levels are neurotoxic ^11, 12^. Ketone bodies are also toxic at elevated levels ^13^. This toxicity at high concentrations of either carbohydrates or ketones underscores the importance of understanding how these fuels interact in the brain to maintain metabolic balance and support mitochondrial function. A key gap in knowledge concerns how acute changes in fuel availability, such as hyperglycemia and ketosis, impact brain metabolism and mitochondrial activity. While studies have focused on isolated effects of glucose or ketones, few have examined their interaction *in vivo*. Given that mitochondrial dysfunction is implicated in various neurological disorders ^14^ and that for nutrients to be considered beneficial, there needs to be a positive effect on mitochondrial function ^15^, understanding the combined effects of glucose and ketones on brain mitochondrial activity is critical to optimizing dietary interventions such as the ketogenic diet.

In this study, we hypothesized that the ketone body BHB can fine-tune mitochondrial function in the brain, acutely mitigating effects associated with stressful metabolic conditions such as acute hyperglycemia or acute brain injury. To test this hypothesis, we developed a novel approach that combines *in vivo* live imaging with an established transgenic reporter mouse model ^16^. This model incorporates an astrocyte-specific promoter for glial fibrillary acidic protein (GFAP) along with an ATP-dependent luciferase enzyme, enabling us to dynamically assess the acute effects of glucose and BHB on brain mitochondrial activity. Luciferase assays have traditionally been used to estimate ATP levels *in vitro* ^17^. However, the novelty of our approach lies in the ability to monitor ATP levels and mitochondrial activity *in vivo,* using bioluminescence as a real-time measure of mitochondrial function. In these assays, luciferase activity depends on ATP hydrolysis, which acts on luciferin to produce bioluminescence. Fluctuations in ATP levels directly translate to changes in bioluminescence, providing an indirect measure of mitochondrial function in response to glucose and BHB. We also confirmed our data using *ex-vivo* tissue biopsies with oxygen consumption measurements. Moreover, to provide functional evidence linking metabolic modulation by BHB to brain health and further emphasize the potential relevance of ketone balance for cerebral resilience, we tested the BHB effect in brain protection from photothrombotic stroke. Our data indicate that the acute effects of metabolic stress or stroke on the brain can be mitigated by BHB.

## Methods

### Animals

All experiments including animal studies, were conducted in accordance with the Guide for the Care and Use of Laboratory Animals and approved by the Institutional Animal Care and Use Committee (IACUC) at the University of Texas Health Science Center at San Antonio. The study complies with the ARRIVE guidelines 2.0 for reporting animal experiments.

### *In vivo* bioluminescence imaging (BLI)

GFAP+GAPDH+ Dual-Glo mice (Jackson lab stock #009638) were fasted 1 day prior to BLI experiments. 3-5 months old, (n = 12 per group including 6 females and 6 males) were injected with 15 mg/kg CycLuc1 (AOBIOUS cat# AOB1117) and anesthetized using 2.5% isoflurane inhalation. After ensuring appropriate anesthesia, mice were imaged using a Xenogen IVIS spectrum live-imaging system, a non-invasive *in vivo* imaging platform that detects and quantifies bioluminescence in real-time. The mice were injected subcutaneously with CycLuc1 in the scruff of the neck, and bioluminescence images were taken every minute for 35 minutes, without any delay.

### CycLuc1 and imaging

The D-luciferin analog (Cycluc1 ^18^), the luciferase substrate, was chosen at a dose of 15 mg/kg, as this concentration was necessary to achieve a minimum detection threshold of 600 counts, ensuring reliable data quantification. At lower CycLuc1 concentrations, bioluminescence signals were too weak for accurate analysis. Imaging was performed using both an open filter and a 620 nm filter, with a 60-second exposure time, Field of View C (FOV:C), and binning of 8. Signal quantification was performed using fixed-size square regions of interest (ROIs) throughout the imaging session.

### Treatment groups

After 4 minutes of Cycluc1 imaging, mice were given a subcutaneous injection of either 2.52g/kg BHB (Sigma cat# H6501), 2 g/kg glucose (a dose that induces acute hyperglycemia ^19^), will be referred here as high glucose (HG), a combination of both 2.52 g/kg BHB and 2 g/kg glucose (HG + BHB), or control (saline: 1.116mg/kg NaCl) in the scruff of the neck using a 31-gauge BD syringe (BD cat# 328468). The time frame of subcutaneous injection is 10 seconds per mouse with three mice imaged simultaneously.

The same cohort of mice was used for all treatment groups, to adhere to the minimalistic animal sacrifice approach, while allowing for within-mouse comparisons, reducing variability, and enhancing statistical power. All 12 mice received the same treatment sequence: saline, BHB, HG, and HG+BHB, with each experiment set 3-7 days apart. Body weight was measured before each injection to ensure accurate dosing and to monitor the health of the mice (**Supplemental Figure S1 A**). For consistency, the master mix was prepared on the same day for all mice before each injection.

The timing of the glucose, BHB, or saline injections (4 minutes after CycLuc1 injection) was carefully chosen to allow sufficient time for CycLuc1 to establish its biological effect, thereby ensuring reliable bioluminescence readings.

### Blood glucose and ketone levels

Blood glucose and ketone levels were measured using the GK+ Blood Glucose & Ketone Meter Kit (Keto Mojo). A small blood sample was collected via tail vein puncture from each mouse at two timepoints: immediately before injection (0 min) and 30 minutes post-injection. The blood sample was applied to test strips, and concentrations of glucose and ketones were determined using the meter’s electrochemical sensor. Blood glucose and ketone levels were measured 30 minutes after injections to capture peak systemic concentrations of the administered compounds.

### Photothrombotic stroke model

3-5-month-old C57BL/6 mice were obtained from Jackson Laboratories (#000664) and maintained in standard Association for Assessment and Accreditation of Laboratory Animal Care International (AAALAC)-approved animal care facility with controlled temperature and humidity, a 12-hour light/dark cycle and ad libitum access to water and chow. The photothrombotic stroke method was previously described ^20^. In brief, mice were anesthetized with isoflurane. Hair was removed from the top of the head using a chemical depilatory. The mouse was placed on the surgical platform where the head was cleaned.

Mice were given a 5 mg/kg subcutaneous injection of Meloxicam prior to surgery. The mice were injected with a Rose Bengal solution (4 mg/mL injectable saline) by intraperitoneal administration 30 minutes prior to inducing stroke. After 15 minutes of Rose Bengal injection, NaCl control (1.116mg/kg), BHB (2.52 g/kg), HG (2 g/kg), or both [BHB (2.52 g/kg), HG (2 g/kg)] were injected intraperitoneally 15 minutes before the photothrombotic stroke. The mouse was placed on a stereotaxic frame, which supplied isoflurane and oxygen throughout the procedure.

Using aseptic techniques, a 1.5 cm midline incision was made through the skin. The connective tissue covering the skull was removed using small scissors, followed by cleaning the skull surface with a hydrogen peroxide swab. The Bregma was located under the microscope.

The fiber optic pipette probe was placed 1.7 mm lateral to the midline in the right hemisphere and 10 mm from the surface of the skull. The right hemisphere was illuminated through the intact skull with a 561-nm laser fiber optic illuminator (Coherent Sapphire) with a 3-mm beam diameter for 15 minutes. The laser setting was 45 mW at 38%, and all laser procedures were performed in a class 4 laser safety room with a laser curtain, in compliance with applicable safety regulations. After 15 minutes of illumination, the probe was removed, and the incision was closed with sutures and gently cleaned with chlorhexidine. A thin layer of a first-aid antibiotic with pain relief was applied. Mice were injected with 10 mg/kg of Enrofloxacin to treat systemic bacterial infections. After surgery, mice were kept in a recovery chamber (∼37°C) for approximately 1.5 hours and then returned to their cages.

### TTC staining and lesion volume quantification

2,3,5-Triphenyltetrazolium chloride (TTC, Sigma Aldrich, # T8877) staining was performed as previously described ^20^. Briefly, 24 hours after stroke brains were removed and placed in ice-cold PBS for 5 min on ice. Then brains were placed in a tissue matrix (Ted Pella, #15050), sliced into 1 mm thick coronal sections through the brain, and placed in solution with TTC for 16 min at 37°C, turning over halfway through incubation. Once stained, sections were scanned.

### Scanner-based image acquisition and infarct size determinations

TTC-stained coronal sections were imaged using an Epson V850 Pro scanner set to professional mode (24-bit color, 1,200 dpi, no color correction, gamma 2.2), with slices placed between transparencies to avoid damage (ordered from rostral to caudal). The scanner settings included medium exposure and a 1 × 1 pixel densitometer sampling area. A 15 cm ruler was included for pixel-to-distance calibration, which was performed in NIH ImageJ (v1.53K)^21^, using a horizontal line on the ruler to set the known distance (10 mm = 235.0021 pixels) and a pixel aspect ratio of 1.0. Lesion areas were outlined with the freehand tool after zooming images to 300%. The lesion volume for each slice was calculated as the average of the front and back side areas multiplied by the slice thickness (1 mm). Total infarct volumes were obtained by summing slice volumes for each mouse brain. Detailed method can be found in our previous study ^20^.

### Seahorse analysis

Brain tissue samples were obtained as 1 mm biopsies from experimental animals after euthanasia for *ex vivo* analysis of oxygen consumption rate (OCR) using the Seahorse XF Analyzer (Agilent). Basal OCR measurements were recorded for each group under various conditions, including incubation with normal glucose (NG, 5.5 mM) alone as control, high glucose (HG, 25 mM) alone, and BHB (BHB, 20 mM). Combination treatments of NG and BHB, as well as HG and BHB, were tested. The metabolic modulators were sequentially injected in the following order: Oligomycin (Olig) (1 μM) (Sigma), ATP synthase inhibitor was used to assess ATP-Linked respiration by measuring the decrease in OCR following its addition. FCCP (fluoro-carbonyl cyanide phenylhydrazone, 1 μM), a mitochondrial uncoupler, was added to collapse the proton gradient and measure maximal respiration, representing the full capacity of the electron transport chain. AA/ROT (antimycin A and rotenone, 1 μM each), inhibitors of complex III and complex I, respectively, were used to block mitochondrial respiration completely.

These sequential injections allowed the assessment of key mitochondrial parameters: Basal respiration, ATP production, proton leak, coupling efficiency, maximal respiration, spare respiratory capacity (SRC), non-mitochondrial OCR, and acute OCR response reflecting the immediate mitochondrial response to substrate addition. This protocol is described in Agilent Seahorse XF Cell Mito Stress Test user guide and in our previous study ^22^.

### Statistical analysis

Statistical analysis was performed using GraphPad Prism 10. For comparisons between 2 groups with repeated measurements a paired two-tailed t-test was used. For multiple comparisons involving repeated measures, a repeated measures one-way ANOVA followed by Tukey’s post hoc test was applied. For comparisons between more than 2 independent groups, a one-way ANOVA followed by Tukey’s post hoc test was applied. The significance level was set at p < 0.05. Graphs display group means ± standard deviation (SD).

## Results

### Acute hyperglycemia decreases ATP flux in the mice brain

The adult brain is widely recognized as a glucose metabolizing organ ^23^. To assess the effect of acute hyperglycemia on brain metabolism, we employed bioluminescence imaging experiments to measure ATP-dependent luciferase activity in transgenic Dual Glo mice expressing luciferase in astrocytes (**Figure 1 A)**. Representative images of mice subsequently injected with either NaCl control (1.16 mg/kg) or HG (2g/kg) are shown in **Figure 1 B**. Note the decrease in photon flux in the brain region after HG injections. We found the bioluminescent signals (photons per second) were consistently lower in mice injected with HG compared to control mice (**Figure 1 C**). The peak total photon flux occurred approximately 10 minutes post-injection for both HG-injected and vehicle-injected mice, at 6.86 x 10^6^ ± 0.75 x 10^6^ (n = 12) and 8.44 x 10^6^ ± 0.65 x 10^6^ (n = 12), respectively. For comparison with tissue responses outside the brain, we simultaneously monitored photon flux in the foot region. We found no statistical differences in the total flux emitted from feet in mice 10 minutes post-injection with either HG (1.96 x 10^6^ ± 0.23 x 10^6^, n=12) or saline (2.26 x 10^6^ ± 0.32 x 10^6^, n=12) (**Supplemental Figure S1 B**). The data indicates a reduction in brain ATP generation with high glucose.

**Figure 1.**
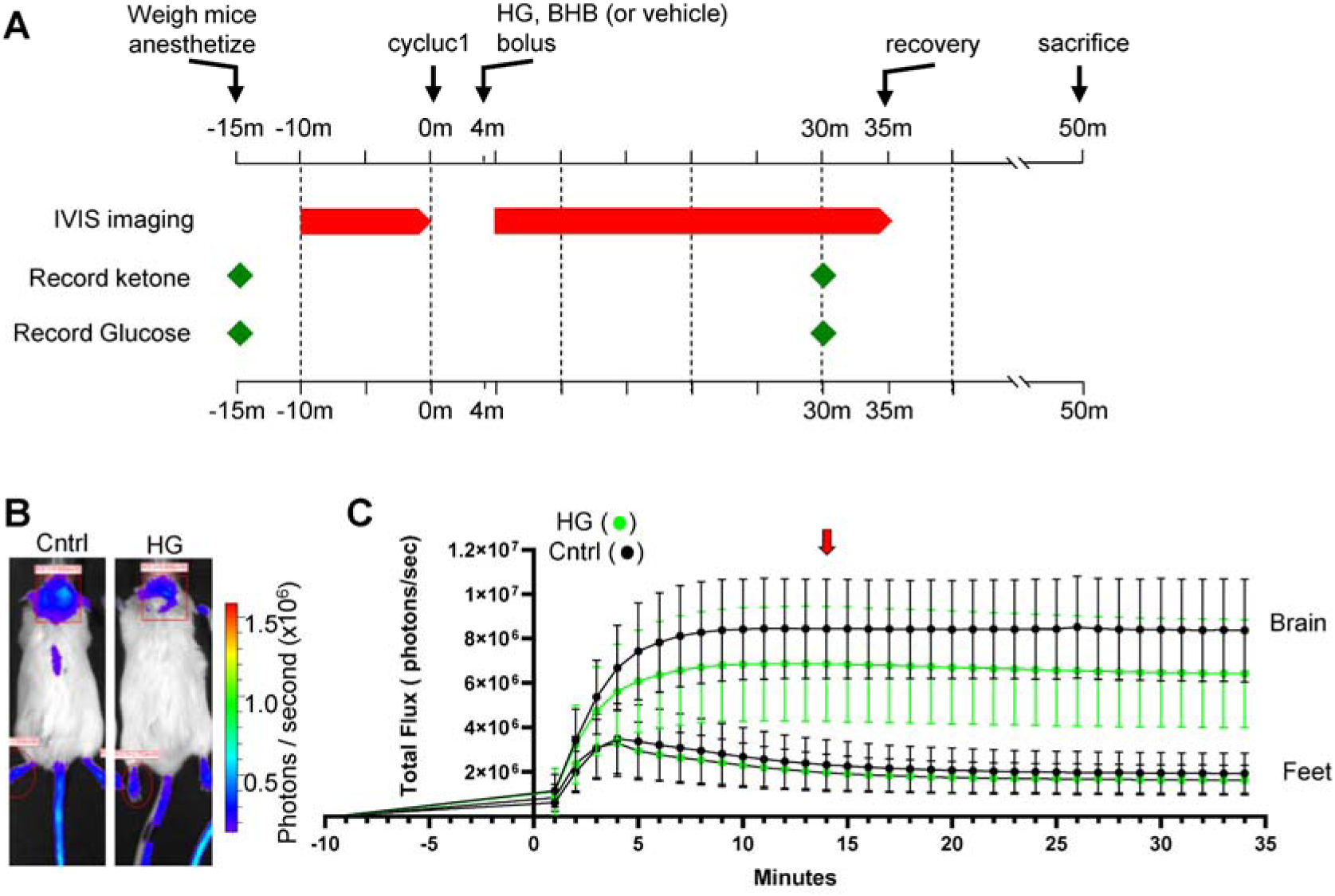
Luciferin-luciferase bioluminescence decreases in response to subcutaneous injections of high glucose (HG). (**A**) Experimental timeline of luciferin-luciferase bioluminescence imaging experiments. (**B**) Transgenic mice expressing luciferase enzyme in astrocytes (Gfap promoter), 14 minutes after injection of luciferin substrate and 10 minutes after injection of either NaCl (control) or HG (2g/kg). Region of interest measured for bioluminescence is shown with red rectangle. Minor expression of luciferase in the skin is observed in tail and feet. The regions of interest over mouse feet, indicated by red ovals, were used as control signals from mice injected with either NaCl or HG. (**C**) Graphical representation of the bioluminescence emission expressed as flux (photons/sec) *vs*. time in minutes. The red arrow indicates the time of image acquisition at peak flux time. Graphs represent mean ± SD.

### **β**-hydroxybutyrate (BHB) increases ATP flux in the mice brain

We next measured the impact of injecting the ketone BHB on brain metabolism in Dual Glo mice. As above, adult Dual Glo mice were injected with the D-luciferin analog Cycluc1. Four minutes later, mice were injected with BHB (2.52 g/kg), and images were acquired on the Xenogen IVIS system. We observed a higher level of photon flux in the brain region of BHB-injected mice compared to saline-injected mice, which was visually apparent 10 minutes post-injection (**Figure 2 A**). Minute-by-minutes changes in total photon flux after BHB injections are presented (**Figure 2 B**). For comparison, the total photon flux for each minute has been replotted from Figure 1 for HG (green dashed line) and control mice (black dashed line). The individual photon flux values for each mouse, recorded 10 minutes post-injection of saline, BHB and HG, are presented in **Figure 3 A and B**. Together, these studies suggest that acute BHB treatment may improve ATP flux.

**Figure 2.**
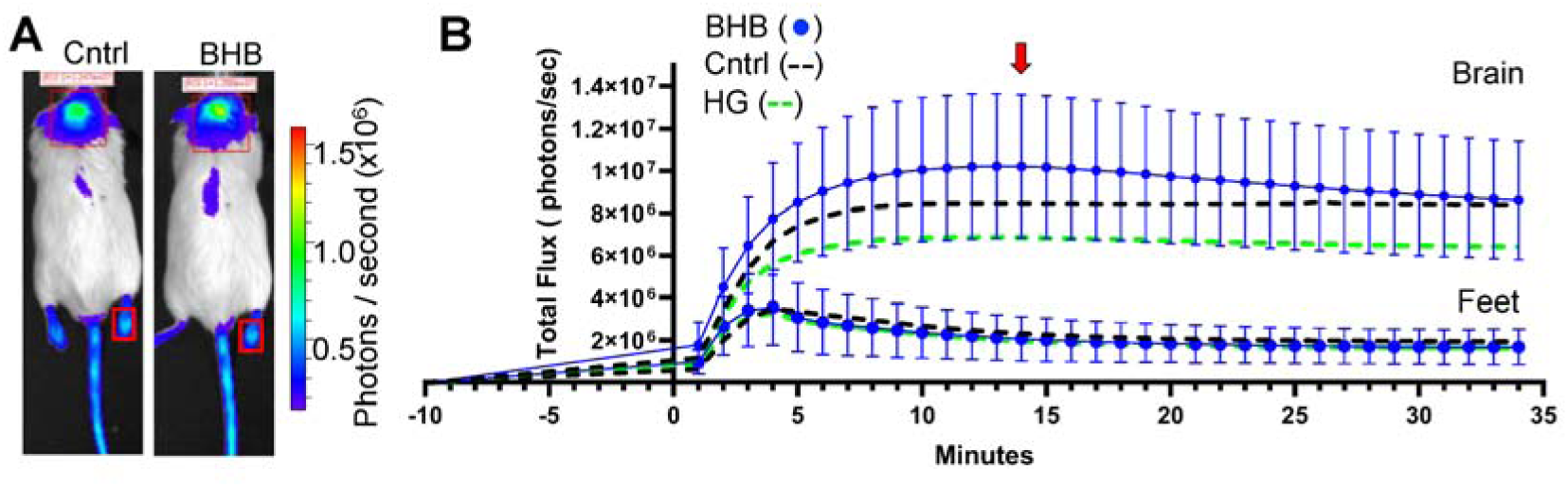
Luciferin-luciferase bioluminescence changes in response to subcutaneous injections of β-hydroxybutyrate (BHB). (**A**) Transgenic mice expressing luciferase enzyme in astrocytes (Gfap promoter), 14 minutes after injection of luciferin substrate or 10 minutes after injection of either NaCl (control) or BHB (2.5 g/kg). Region of interest measured for bioluminescence is shown with red rectangle. Minor expression of luciferase in the skin is observed in tail and feet. The regions of interest over mouse feet, indicated by smaller red rectangles, were used as control signals from mice injected with either NaCl or BHB. (**B**) Graphical representation of the bioluminescence emission expressed as flux (photons/sec) *vs*. time in minutes. The red arrow indicates the time of image acquisition at peak flux time. Graphs represent mean ± SD.

**Figure 3.**
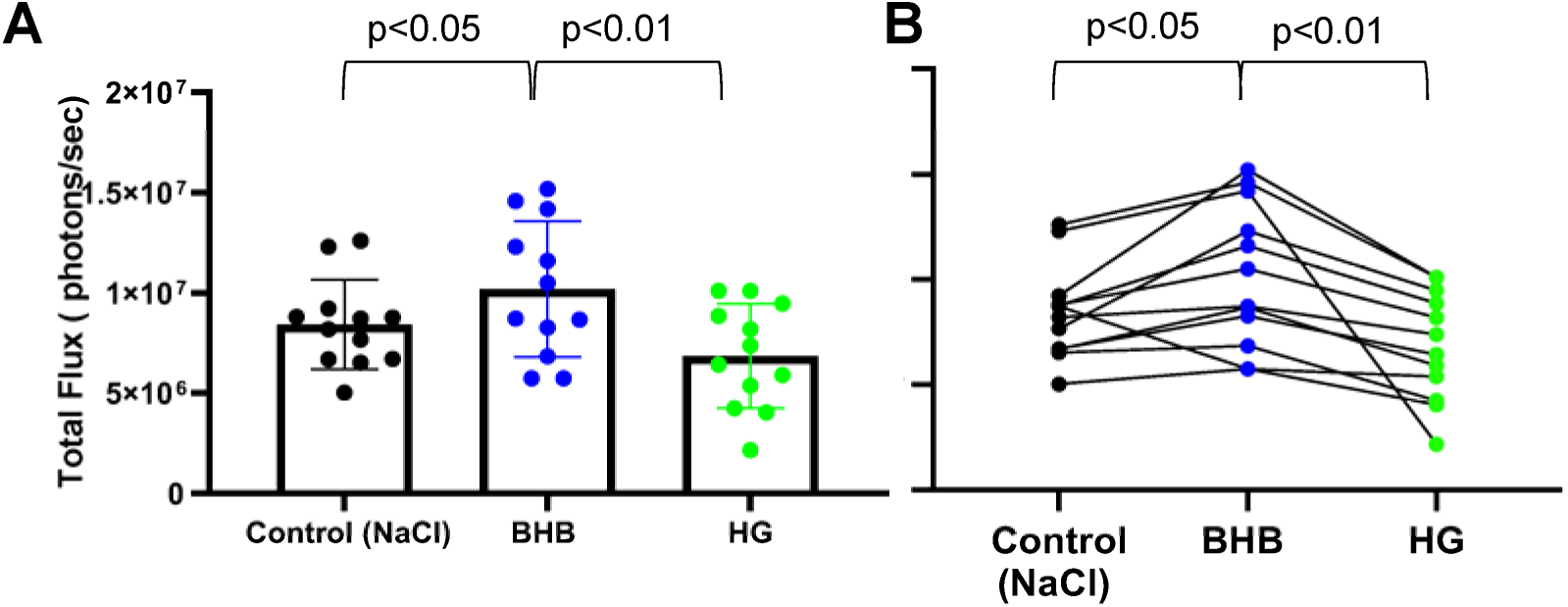
β-hydroxybutyrate (BHB) increases luciferin-luciferase bioluminescence in mouse brain. (**A**) Bar plot comparing bioluminescence in individual mice 14 minutes after injection of luciferin substrate or 10 minutes post-injection of either NaCl Control, BHB (2.52g/kg), or HG (2g/kg). Graph represents mean ± SD. (**B**) Paired data plot showing photon flux measurements for each individual mouse under the three conditions. Each line connects data points corresponding to the same mouse. Each dot represents the photon flux of an individual mouse. Statistical significance between conditions was assessed using repeated measures ANOVA test with Tukey’s multiple comparison test.

### **β**-hydroxybutyrate (BHB) sustains brain metabolism in acute hyperglycemic conditions

We also measured the impact of simultaneously injecting HG and the ketone BHB on brain metabolism. As above, dual glo mice were injected with CycLuc1 followed 4 minutes later by mixed solution of HG and BHB, the peak photon flux for HG and BHB measured in the brain region (red symbols/line, **Figure 4**) was comparable to control mice injected with NaCl control alone (black dashed line, **Figure 4**), albeit with a slower initial rise in photon flux for the first 10 minutes. By minute 14 (equivalent to 10 minutes post-injection of HG+ BHB), photon flux levels were identical to control values. Histogram and paired data plots of brain photon flux for individual mice are presented in **Figure 4 B and C**. No significant differences in photon flux were observed between control and HG + BHB injected mice.

**Figure 4.**
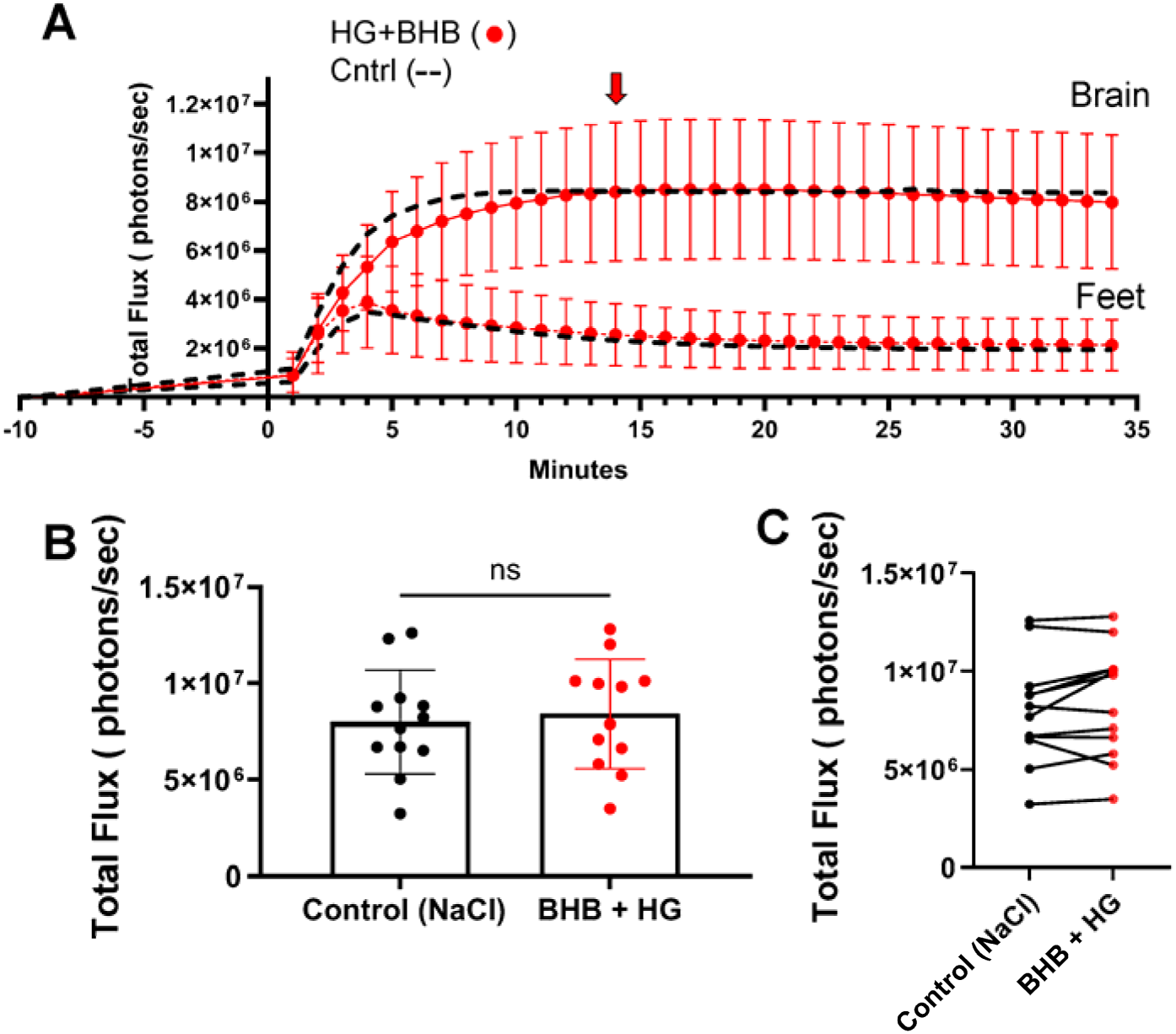
β-hydroxybutyrate (BHB) co-administered with high glucose (HG) maintains bioluminescence at control levels. (**A**) Bioluminescence of mice expressing luciferase under Gfap promoter (astrocytes). Mice were injected with luciferin substrate, followed, 4 minutes, later by either control (NaCl), HG + BHB. Photon flux was measured every minute post-luciferin injection. BHB (2.52g/kg) and HG (2g/kg) were injected together. The red arrow indicates the time of image acquisition at peak flux time. (**B**) Bar plot comparing photon flux at 14 minutes post-luciferin injection between the control (NaCl) group and the HG + BHB group. (**C**) Paired data plot showing individual photon flux measurements for each mouse. Each line connects the photon flux values of the same mouse under both conditions, demonstrating within-subject variability. All measurements were acquired on Xenogen IVIS system. Each dot represents an individual mouse, and graphs represent means ± SD. Paired two-tailed t-test indicated no significant difference (ns).

### **β**-hydroxybutyrate (BHB) ameliorates blood glucose level amid acute hyperglycemia setting

We measured blood levels of HG and BHB in mice, both before and 30 minutes after subcutaneous injections of saline, HG, BHB or HG+BHB. As expected, blood glucose levels were not affected by saline or BHB injections (**Figure 5 A**). However, the average blood glucose significantly increased (p < 0.0001) from 177.6 ± 6.8 mg/dl (n = 9) at 0 minutes to 352.3 ± 13.6 (n = 9), 30 minutes after mice had been injected with HG (**Figure 5 A**). Blood glucose levels were also not significantly changed when HG and BHB were injected together. Saline and HG injections had no effect on blood ketone levels, while BHB injections significantly increased (p < 0.0001) the blood ketone levels to 7.54 ± 0.23 mM (n = 9) 30 minutes post-injection (**Figure 5 B**). When BHB was co-injected with HG, ketone levels rose to 4.61 ± 0.38 mM (n = 9).

**Figure 5.**
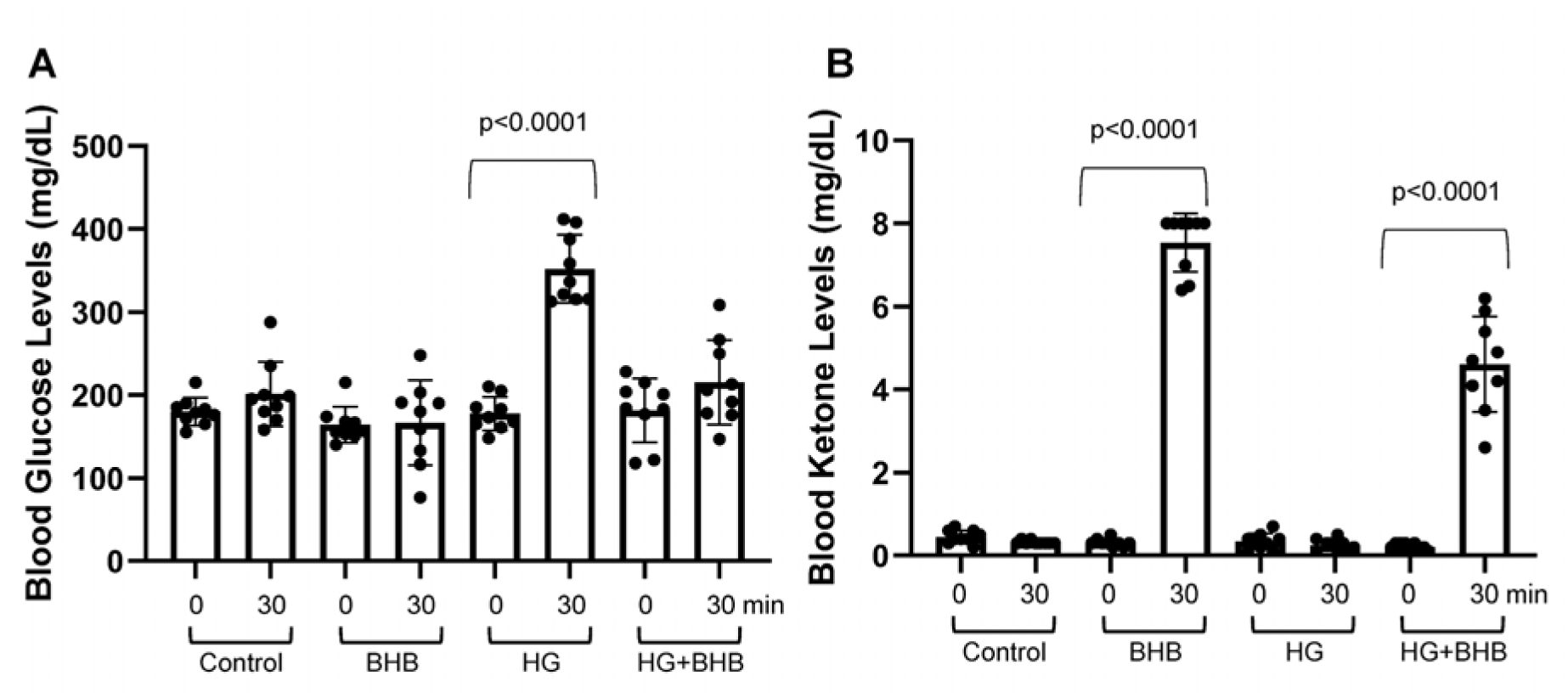
Blood glucose and ketone concentrations after subcutaneous injection of high glucose (HG) and β-hydroxybutyrate (BHB). (**A**) Blood glucose concentrations in mice before (0min) and after (30min) injections of NaCl control, BHB (2.52g/kg), HG (2g/kg) and [BHB (2.52g/kg) +HG (2g/kg)] together as indicated. (**B**) Blood ketone concentrations in mice before (0min) and after (30min) injections of NaCl control, BHB (2.52g/kg), HG (2g/kg) and [BHB (2.52g/kg) +HG (2g/kg)] as indicated. Each symbol is an individual mouse. Data are presented as mean ± SD. Statistical significance was determined using Tukey’s multiple comparison test.

### Effects of **β**-hydroxybutyrate (BHB) on mitochondrial respiration parameters under normoglycemic and hyperglycemic conditions

Oxygen consumption rate (OCR) measurements of *ex vivo* brain tissue (1 mm biopsies), which indicate mitochondrial activity, are shown in **Figure 6 A.** OCR dynamics over time for all treatments (NG, NG+BHB, HG, HG+BHB) are illustrate in **Figure 6 B,** highlighting the mitochondrial response to the sequential addition of modulators and substrates. Basal respiration, representing the OCR in the absence of additional substrates or stressors, showed no significant differences between groups. The normal glucose (NG) control group exhibited a basal OCR of 89.25 pmol O_₂_/min, compared to 71.77 pmol O_₂_/min in the NG+BHB group, 74.3 pmol O2/min in the HG group and 61.14 pmol O_₂_/min in the HG+BHB group (**Figure 6 C**). These findings suggest that BHB does not alter basal mitochondrial activity.

**Figure 6.**
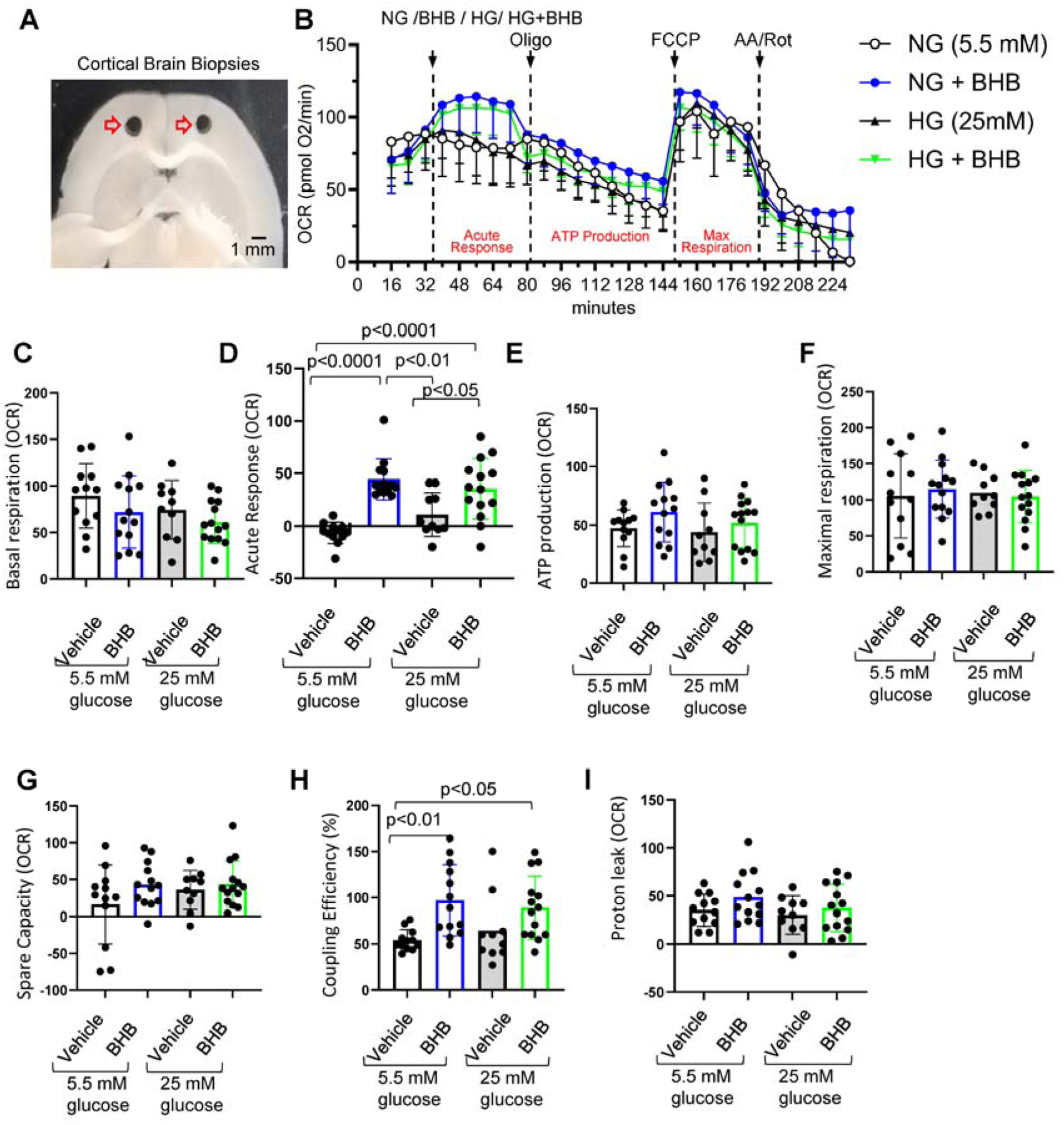
Cortical brain biopsy respiration analysis under normoglycemic and hyperglycemic conditions with and without β-hydroxybutyrate (BHB). (**A**) Microscopic images of cortical mouse brain biopsies retrieved after sacrifice and used for *ex vivo* Seahorse analysis. (**B**) Oxygen consumption rate (OCR), measured in pmol O_₂_/min, plot showing dynamic changes in oxygen consumption across treatments, including normal glucose (NG, 5.5 mM glucose), NG + 20 mM BHB, high glucose (HG, 25 mM glucose), and HG + BHB. OCR values were recorded for 232 minutes, capturing baseline respiration, sequential injection responses, and acute OCR changes following substrate administration. (**C**) Basal respiration, calculated as the last OCR measurement before the acute injection (measurement # 3) minus non-mitochondrial respiration (minimum rate after AA/Rot injection). (**D**) Acute response, calculated as the difference between the fourth OCR measurement after substrate injection (measurement #7) and the last basal OCR measurement before acute injection (measurement #3). (**E**) ATP production, calculated as the difference between last OCR measurement before oligo injection (measurement # 9) and minimum rate measurement after oligo injection (measurement # 18). (**F**) Maximal respiration, calculated as the maximal OCR after FCCP injection (measurement # 20) minus non-mitochondrial respiration. (**G**) Spare respiratory capacity (SRC), calculated as maximal respiration minus basal respiration (**H**) Coupling efficiency, calculated as ATP production/Basal respiration ×100. (**I**) Proton leak, calculated as the minimum OCR after oligo injection (measurement # 18) minus non-mitochondrial respiration. Data was acquired on a Seahorse XF analyzer (Agilent). 24 well system from 4 mice, replicates counts are as such: 12 NG, 10 HG, 13 NG+BHB and 14 HG+BHB. Data are presented as means ± SD. Statistical significance was determined using One-way ANOVA test followed by Tukey’s multiple comparison test. AA/Rot: antimycin A and rotenone; FCCP: carbonyl cyanide-4 (trifluoromethoxy) phenylhydrazone; Oligomycin (Oligo).

Acute OCR response, indicating the immediate reaction of mitochondria to the addition of modulators, showed significant increases in NG+BHB (44.46 pmol O_₂_/min) and HG+BHB (35.57 pmol O_₂_/min) groups compared to NG (–6.5 pmol O_₂_/min) and HG (10.9 pmol O_₂_/min) controls (p < 0.05) (**Figure 6 D)**.

ATP production, indicating mitochondrial efficiency in converting substrates into usable energy, showed a trend toward an increase with BHB in NG (61.15 pmol O_₂_/min) compared to NG controls (47.17 pmol O_₂_/min), though the difference was not statistically significant. In HG conditions, ATP production remained consistent between HG (43.7 pmol O_₂_/min) and HG+BHB (51.79 pmol O_₂_/min) groups (**Figure 6 E)**.

Maximal respiration, which measures the full capacity of the electron transport chain under uncoupled conditions, showed no significant differences between groups (**Figure 6 F).** Spare respiratory capacity (SRC), indicating the difference between maximal respiration and basal respiration and reflecting the ability of mitochondria to respond to increased energy demands, also showed no significant differences across groups (**Figure 6 G).** However, the coupling efficiency, indicating the proportion of oxygen consumption used for ATP production relative to total oxygen consumption, was significantly improved with BHB treatment. Under NG conditions, coupling efficiency increased from 54.1% in controls to 96.95% with BHB, and under HG conditions, coupling efficiency increased from 64.47% to 89.11% (**Figure 6 H**).

Proton leak, representing uncoupled respiration, increased (but not significantly) with BHB treatment in both NG and HG conditions. The NG+BHB group exhibited a higher proton leak compared to controls, suggesting that BHB induces a mild stress adaptation or regulatory adjustment in mitochondrial activity (**Figure 6 I)**. These findings indicate that BHB enhances mitochondrial efficiency and acute responsiveness without altering basal or maximal respiration.

### Acute **β**-hydroxybutyrate (BHB) reduced infarct volume in a thrombotic stroke model

In order to establish a functional link between metabolism and brain health, we investigated the acute effects of acute BHB in a photothrombotic stroke model, measuring infarct volume as a functional outcome (**Figure 7 A).** Mice treated with BHB demonstrated a significant reduction in infarct volume compared to controls. The average infarct volume was 29 ± 2.1 mm³ in the control (NaCl) group, which was reduced to 18 ± 1.9 mm³ in the BHB group (p < 0.001). Mice treated with HG alone had an infarct volume of 24 ± 1.5 mm³, while the combination of HG and BHB resulted in a smaller infarct volume of 20 ± 3.1 mm³ (p = 0.05 compared to NaCl control group) (**Figure 7 B)**. These results provide evidence of a functional acute benefit of BHB, demonstrating its cerebroprotective effects both under normal and hyperglycemic conditions.

**Figure 7.**
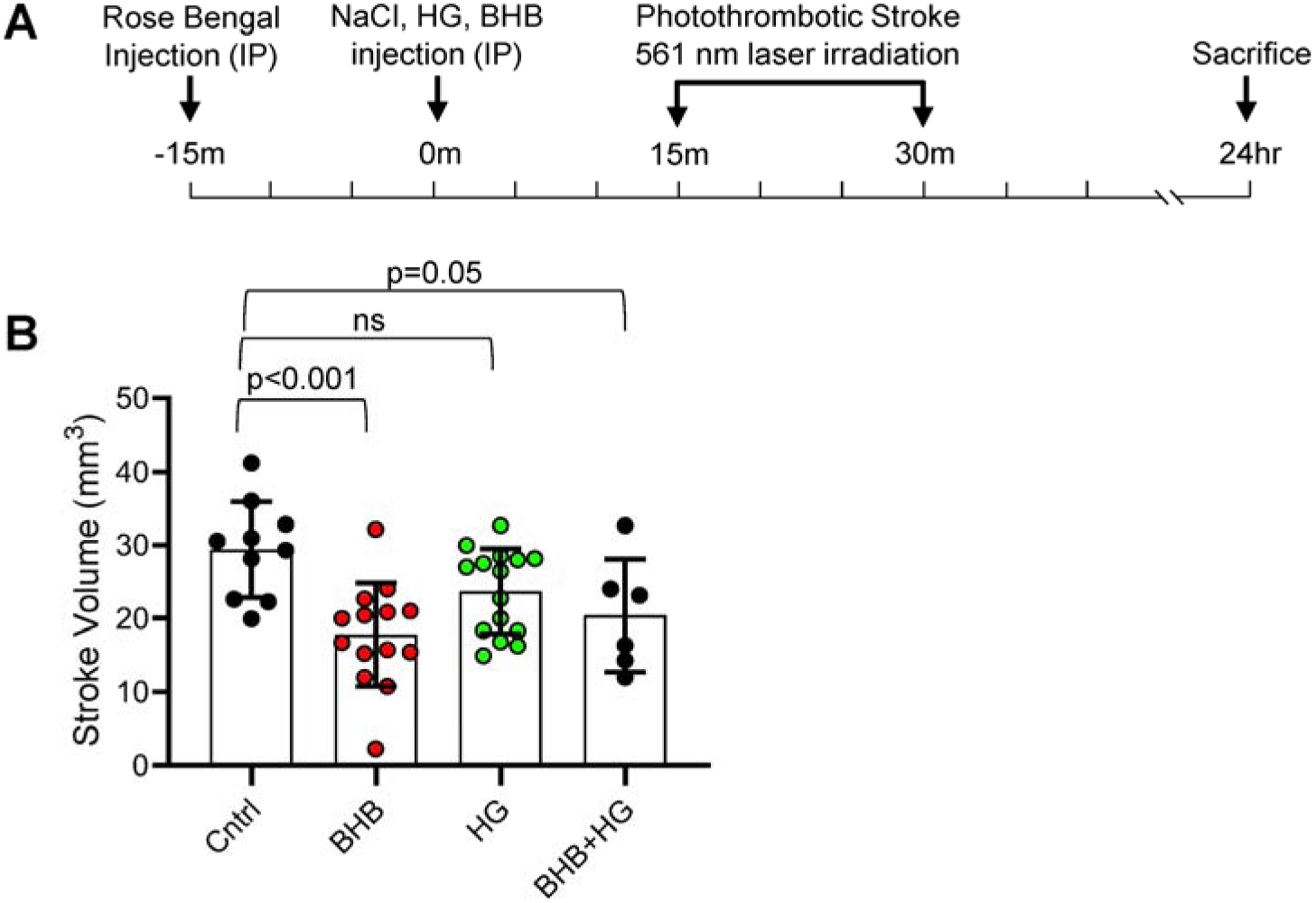
Effect of β-hydroxybutyrate (BHB) on brain stroke infarct volume. (**A**) Schematic of the experimental design. Mice were injected with Rose Bengal solution 30 minutes before stroke induction. Treatments [NaCl (n=10), BHB (n=14), HG (n=15), or HG + BHB (n=6)] were administered intraperitoneally 15 minutes before stroke induction. The photothrombotic stroke was induced using a 561 nm laser for 15 minutes, and mice were sacrificed 45 minutes post-stroke. (**B**) Infarct volumes were measured 1 hour after treatment. Data are presented as means ± SD. Statistical significance was determined using one-way ANOVA test followed by Tukey’s multiple comparison test.

## Discussion

The major goal of our multi-disciplinary study was to identify a mechanism by which a ketogenic diet may be beneficial to the brain. Our hypothesis was that ketone bodies can mitigate acute metabolic stress on the brain. The data in mice presented above indicate that acute hyperglycemia induces a decrease in ATP levels within the brain, while BHB supplementation exhibits promising effects on brain metabolism under such conditions. Specifically, we found that BHB increases ATP flux in the brain and ameliorates both brain metabolism and blood glucose levels during acute hyperglycemia.

BHB may also enhance glucose utilization and insulin sensitivity in non-diabetic individuals, as evidenced by studies such as Myette-Côté et al. ^24^, which demonstrated an improved insulin sensitivity index and reduced glucose area under the curve following BHB supplementation. This highlights the possible ability of BHB to complement insulin action by promoting glucose regulation and metabolic balance.

We also investigated the impact of BHB on mitochondrial function using OCR analysis. We found that BHB significantly enhanced coupling efficiency in both normoglycemic and hyperglycemic conditions, indicating improved mitochondrial efficiency, possibly by decreasing proton leak and thereby optimizing oxygen utilization for ATP production. This increase in coupling efficiency was not accompanied by a significant change in ATP production, suggesting that BHB improves mitochondrial performance by minimizing inefficiencies (e.g. proton leak or non-productive respiration) rather than directly increasing energy output. This is also consistent with the idea that cellular ATP production is demand-driven and may not rise unless energy requirements increase ^25, 26^. An important caveat of this interpretation is that these measurements were performed on brain biopsies, where the heterogeneous metabolic activity of neurons, astrocytes, and other brain cells ^27^, as well as the high basal metabolic rate of brain tissue ^28^, could mask detectable changes in cell-specific ATP production Additionally, BHB increased the acute OCR response following substrate addition, particularly under HG conditions, suggesting a rapid and adaptive mitochondrial response. This enhanced acute OCR response may indicate that BHB primes mitochondria for metabolic flexibility, enabling a quicker adjustment to substrate availability and metabolic stress. These findings align with prior studies, such as Shimazu et al. ^29^, which demonstrated that BHB enhances energy efficiency and reduces oxidative stress in neuronal models.

The mild and potential increase in proton leak, observed with BHB treatment, may reflect a regulatory mechanism to mitigate reactive oxygen species (ROS) production, possibly by dissipating excess mitochondrial membrane potential, as suggested in other studies ^30–32^. These adaptations suggest an acute role of BHB in improving efficient energy utilization while enhancing mitochondrial resilience even under metabolic stress.

In addition, our photothrombotic stroke data reveal that BHB significantly reduced infarct size, providing direct functional evidence of its cerebroprotective effects.

### Acute hyperglycemia alters brain metabolism even in the absence of diabetes

Energy metabolism in the brain is a complex process that involves the regulation of glucose uptake and utilization, as well as the metabolism of phospholipids. Glucose is a crucial energy substrate for the brain, and its uptake is tightly regulated to meet the high energy demands of the brain ^27, 33^. The brain’s reliance on glucose as a primary energy source is well-established, and alterations in glucose metabolism can have significant effects on brain function ^34^.

Here we highlight that acute hyperglycemia could alter the brain’s metabolic dynamics even in the absence of diabetes. A study in humans employing non-invasive magnetic resonance techniques showed that ingesting 50 g glucose decreased global cerebral metabolic rate of oxygen in a time-dependent pattern, which was accompanied by a reduction in oxygen extraction fraction and metabolism of oxygen ^35^.

Our Seahorse data demonstrated that BHB enhanced mitochondrial oxygen consumption even under hyperglycemic potential stress, as evidenced by improved coupling efficiency and acute OCR response. These mitochondrial adaptations suggest that BHB provides an acute metabolic advantage in balancing energy demands and limiting hyperglycemia-induced inefficiencies.

### BHB is a brain-friendly energy substitute that enhances oxygen consumption rate

BHB has gained attention for its potential as an alternative energy source in the absence of glucose. Studies have shown that BHB not only serves as an energy source but also exhibits neuroprotective properties in models of neurodegenerative diseases like Parkinson’s, Huntington’s, and Alzheimer’s ^36–39^.

Our study is the first to investigate the exogenous BHB effect under normoglycemia and acute hyperglycemia and provides an explanation of how BHB could protect the brain by improving mitochondrial efficiency.

Our data indicate that ATP flux under BHB-treatment is increased and returns to control levels over time, suggesting a transient metabolic boost provided by BHB under acute conditions. Conversely, HG-treatment exhibited a consistent decrease in flux, indicating a sustained negative impact on mitochondrial function. The HG + BHB treatment maintained a control flux level, but this was associated with a slight decrease in ATP flux over time, suggesting the beneficial effects of BHB may be temporary. This observation emphasizes the need for studies exploring prolonged BHB exposure or repeated dosing to determine its potential for sustained metabolic benefits.

BHB’s anti-inflammatory and antioxidant properties may help mitigate oxidative stress and inflammation associated with diabetes, thereby enhancing overall metabolic health and lowering the risk of diabetes-related complications ^39–41^. We observed that BHB increases brain OCR in normal or high glucose conditions; this may be partly explained by the effect of BHB on astrocytic glucose consumption, as shown in Valdebenito et al. (2015) ^42^, where BHB inhibits glucose uptake in astrocytes. Modulation of astrocytic glucose metabolism may limit the immediate impact of high glucose levels in the brain, shifting energy metabolism toward BHB utilization and protecting against hyperglycemia-induced metabolic alterations. Prior studies have demonstrated that 30 mg/kg of BHB administered to mice post brain-ischemia-reperfusion improved mitochondrial function and enhanced OCR ^43^. Interestingly, BHB was found to inhibit apoptosis in astrocytes exposed to high glucose concentration ^44^.

In addition to its metabolic effects, our study demonstrated that BHB significantly reduced infarct size in a photothrombotic stroke model. This model is particularly relevant as it induces localized ischemic injury and replicates the acute metabolic and mitochondrial stress that occurs during stroke ^20, 45^, providing a platform to evaluate the immediate effects of metabolic modulators such as BHB.

Our data are supported by recent studies showing that BHB administration via drinking water for 14 days before stroke improved cognitive and motor recovery post-stroke through activation of the nuclear factor erythroid 2-related factor 2 (Nrf2) / antioxidant response element (ARE) pathway ^46^. Furthermore, post-stroke BHB delivery has been associated with reduced peri-infarct glucose uptake, ROS production and astrogliosis, alongside improved neuronal functioning ^47^. Research has also shown that acute BHB administration immediately after reperfusion in a mouse model of transient ischemia enhanced mitochondrial respiration, particularly through improved complex I and II activity^43^. Interestingly, our results demonstrate that the neuroprotective effect of BHB, even when administered acutely before ischemia, underscores its translational potential in mitigating brain injury during acute metabolic and ischemic stress.

### Exogenous BHB, at moderate concentrations, has positive physiological effects

We and others demonstrate that exogenous BHB treatment regulates blood sugar levels by providing an alternative fuel source for insulin-resistant cells, potentially improving insulin sensitivity and reducing insulin resistance ^48–50^. Exogenous ketosis achieved without carbohydrate restriction has been shown to reduce blood glucose levels effectively. For instance, infusion of BHB has been observed to lower blood glucose levels in both animals ^50–53^ and human subjects ^54^. However, it is noteworthy that BHB levels increase in correlation with the severity of diabetic ketoacidosis ^55^. The consideration of BHB’s safe-dose administration is critical to procure brain and systemic safety.

Hence, it is important to appreciate that a low concentration of exogenous BHB administered to mice has a significant protective effect on the brain under both normal and acute hyperglycemic conditions. This safe, low-level BHB administration is reiterated with other studies demonstrating the potential reversal of type 2 diabetes through very low carbohydrate or ketogenic diets ^50^. Thus, monitoring ketone levels during a ketogenic diet is crucial to ensure the diet’s effectiveness and safety.

**Limitations** include a lack of direct measures of ATP and that only astrocytes were assessed for the impact of hyperglycemia. In future studies we plan to perform spatial metabolomics to discern ATP, adenosine diphosphate (ADP) and adenosine monophosphate (AMP) *in situ* and co-localize with specific cell types.

Additionally, our findings reflect acute events, which may differ under chronic conditions. Chronic metabolic dysregulation, such as that seen in diabetes, involves long-term adaptations in glucose transporters and mitochondrial function, which could impact BHB’s metabolism and efficacy. Future studies extending the duration of experiments or employing models of chronic hyperglycemia are necessary to explore the translational relevance of our findings to diseased states.

**In conclusion,** our study provides a firm foundation for future clinical studies that investigate BHB as a variable in determining brain metabolism protection especially in the setting of a ketogenic diet that is mixed with low or high carbohydrate content.

## Author contributions

KS and JDL conceived the project, designed, and provided resources and funding acquisition; DMH, AS, DL, and JW designed, performed experiments, analyzed, and interpreted data; AS, DMH, KS, and JDL wrote the manuscript. DMH and AS contributed equally. JDL provided oversight of the project.

## Statements and Declarations

### Ethical considerations

This study does not involve human research. All experiments including animal studies, were conducted in accordance with the Guide for the Care and Use of Laboratory Animals and approved by the Institutional Animal Care and Use Committee (IACUC) at the University of Texas Health Science Center at San Antonio.

### Consent to participate

Not applicable. This study does not involve human research.

### Consent for publication

Not applicable. This study does not involve human research.

### Declaration of conflicting interest

The funder, Astrocyte Pharmaceuticals, reviewed and approved the manuscript. JDL is co-founder and advisor for Astrocyte Pharmaceuticals. JDL is also an equity holder in Astrocyte. All other authors declare no competing interests or personal relationships that could influence the work reported in this paper.

### Funding statement

This study received funding from Astrocyte Pharmaceuticals, Inc. (JDL) and the 80/20 Foundation (KS). AS was supported by the National Center for Advancing Translational Sciences, National Institutes of Health (NIH), through Grant TL1 TR002647, and by the National Heart, Lung, and Blood Institute, NIH, through Grant T32 HL007446. The content is solely the responsibility of the authors and does not necessarily represent the official views of the NIH. Images were generated in the Core Optical Imaging Facility which is supported by UT Health San Antonio and NIH-NCI P30 CA54174.

### Data availability

All data generated or analyzed during this study are included in the figures and supplementary materials of this manuscript. Additional clarification or access to raw data are available upon reasonable request from the corresponding author.

## References

1. Baranano KW and Hartman AL. The ketogenic diet: uses in epilepsy and other neurologic illnesses. Curr Treat Options Neurol 2008; 10: 410–419. DOI: 10.1007/s11940-008-0043-8.

2. Bueno NB, de Melo IS, de Oliveira SL, et al. Very-low-carbohydrate ketogenic diet v. low-fat diet for long-term weight loss: a meta-analysis of randomised controlled trials. Br J Nutr 2013; 110: 1178–1187. 20130507. DOI: 10.1017/S0007114513000548.

3. Dynka D, Kowalcze K, Charuta A, et al. The Ketogenic Diet and Cardiovascular Diseases. Nutrients 2023; 15 20230728. DOI: 10.3390/nu15153368.

4. Bhanpuri NH, Hallberg SJ, Williams PT, et al. Cardiovascular disease risk factor responses to a type 2 diabetes care model including nutritional ketosis induced by sustained carbohydrate restriction at 1 year: an open label, non-randomized, controlled study. Cardiovasc Diabetol 2018; 17: 56. 20180501. DOI: 10.1186/s12933-018-0698-8.

5. Gjuladin-Hellon T, Davies IG, Penson P, et al. Effects of carbohydrate-restricted diets on low-density lipoprotein cholesterol levels in overweight and obese adults: a systematic review and meta-analysis. Nutr Rev 2019; 77: 161–180. DOI: 10.1093/nutrit/nuy049.

6. Brinkworth GD, Buckley JD, Noakes M, et al. Long-term effects of a very low-carbohydrate diet and a low-fat diet on mood and cognitive function. Arch Intern Med 2009; 169: 1873–1880. DOI: 10.1001/archinternmed.2009.329.

7. Altayyar M, Nasser JA, Thomopoulos D, et al. The Implication of Physiological Ketosis on The Cognitive Brain: A Narrative Review. Nutrients 2022; 14 20220125. DOI: 10.3390/nu14030513.

8. Holsen LM, Hoge WS, Lennerz BS, et al. Diets Varying in Carbohydrate Content Differentially Alter Brain Activity in Homeostatic and Reward Regions in Adults. J Nutr 2021; 151: 2465–2476. DOI: 10.1093/jn/nxab090.

9. Wei SJ, Schell JR, Chocron ES, et al. Ketogenic diet induces p53-dependent cellular senescence in multiple organs. Sci Adv 2024; 10: eado1463. 20240517. DOI: 10.1126/sciadv.ado1463.

10. Kawahito S, Kitahata H and Oshita S. Problems associated with glucose toxicity: role of hyperglycemia-induced oxidative stress. World J Gastroenterol 2009; 15: 4137–4142. DOI: 10.3748/wjg.15.4137.

11. Quincozes-Santos A, Bobermin LD, de Assis AM, et al. Fluctuations in glucose levels induce glial toxicity with glutamatergic, oxidative and inflammatory implications. Biochim Biophys Acta Mol Basis Dis 2017; 1863: 1–14. 20160920. DOI: 10.1016/j.bbadis.2016.09.013.

12. Bahniwal M, Little JP and Klegeris A. High Glucose Enhances Neurotoxicity and Inflammatory Cytokine Secretion by Stimulated Human Astrocytes. Curr Alzheimer Res 2017; 14: 731–741. DOI: 10.2174/1567205014666170117104053.

13. Fedorovich SV, Voronina PP and Waseem TV. Ketogenic diet versus ketoacidosis: what determines the influence of ketone bodies on neurons? Neural Regen Res 2018; 13: 2060–2063. DOI: 10.4103/1673-5374.241442.

14. Alshial EE, Abdulghaney MI, Wadan AS, et al. Mitochondrial dysfunction and neurological disorders: A narrative review and treatment overview. Life Sci 2023; 334: 122257. 20231108. DOI: 10.1016/j.lfs.2023.122257.

15. Rodriguez-Cano AM, Calzada-Mendoza CC, Estrada-Gutierrez G, et al. Nutrients, Mitochondrial Function, and Perinatal Health. Nutrients 2020; 12 20200721. DOI: 10.3390/nu12072166.

16. Cho W, Hagemann TL, Johnson DA, et al. Dual transgenic reporter mice as a tool for monitoring expression of glial fibrillary acidic protein. J Neurochem 2009; 110: 343–351. 20090505. DOI: 10.1111/j.1471-4159.2009.06146.x.

17. Turman MA and Mathews A. A simple luciferase assay to measure atp levels in small numbers of cells using a fluorescent plate reader. In Vitro Cell Dev Biol Anim 1996; 32: 1–4. DOI: 10.1007/BF02722985.

18. Evans MS, Chaurette JP, Adams ST, Jr., et al. A synthetic luciferin improves bioluminescence imaging in live mice. Nat Methods 2014; 11: 393–395. 20140209. DOI: 10.1038/nmeth.2839.

19. Hu J, Zhong X, Yang X, et al. A novel inducible acute hyperglycemia mouse model for assessing 6-KTP. Biomed Rep 2019; 11: 110–114. 20190708. DOI: 10.3892/br.2019.1228.

20. Fisher ES, Chen Y, Sifuentes MM, et al. Adenosine A1R/A3R agonist AST-004 reduces brain infarction in mouse and rat models of acute ischemic stroke. Front Stroke 2022; 1 20221122. DOI: 10.3389/fstro.2022.1010928.

21. Schneider CA, Rasband WS and Eliceiri KW. NIH Image to ImageJ: 25 years of image analysis. Nat Methods 2012; 9: 671–675. DOI: 10.1038/nmeth.2089.

22. Darshi M, Tumova J, Saliba A, et al. Crabtree effect in kidney proximal tubule cells via late-stage glycolytic intermediates. iScience 2023; 26: 106462. 20230321. DOI: 10.1016/j.isci.2023.106462.

23. Mergenthaler P, Lindauer U, Dienel GA, et al. Sugar for the brain: the role of glucose in physiological and pathological brain function. Trends Neurosci 2013; 36: 587–597. 20130820. DOI: 10.1016/j.tins.2013.07.001.

24. Myette-Cote E, Neudorf H, Rafiei H, et al. Prior ingestion of exogenous ketone monoester attenuates the glycaemic response to an oral glucose tolerance test in healthy young individuals. J Physiol 2018; 596: 1385–1395. 20180302. DOI: 10.1113/JP275709.

25. Rolfe DF and Brown GC. Cellular energy utilization and molecular origin of standard metabolic rate in mammals. Physiol Rev 1997; 77: 731–758. DOI: 10.1152/physrev.1997.77.3.731.

26. Brand MD, Chien LF and Diolez P. Experimental discrimination between proton leak and redox slip during mitochondrial electron transport. Biochem J 1994; 297 (Pt 1): 27–29. DOI: 10.1042/bj2970027.

27. Belanger M, Allaman I and Magistretti PJ. Brain energy metabolism: focus on astrocyte-neuron metabolic cooperation. Cell Metab 2011; 14: 724–738. DOI: 10.1016/j.cmet.2011.08.016.

28. Shulman RG, Rothman DL, Behar KL, et al. Energetic basis of brain activity: implications for neuroimaging. Trends Neurosci 2004; 27: 489–495. DOI: 10.1016/j.tins.2004.06.005.

29. Shimazu T, Hirschey MD, Newman J, et al. Suppression of oxidative stress by beta-hydroxybutyrate, an endogenous histone deacetylase inhibitor. Science 2013; 339: 211–214. 20121206. DOI: 10.1126/science.1227166.

30. Nanayakkara GK, Wang H and Yang X. Proton leak regulates mitochondrial reactive oxygen species generation in endothelial cell activation and inflammation – A novel concept. Arch Biochem Biophys 2019; 662: 68–74. 20181203. DOI: 10.1016/j.abb.2018.12.002.

31. Murphy MP. How mitochondria produce reactive oxygen species. Biochem J 2009; 417: 1–13. DOI: 10.1042/BJ20081386.

32. Zorov DB, Juhaszova M and Sollott SJ. Mitochondrial reactive oxygen species (ROS) and ROS-induced ROS release. Physiol Rev 2014; 94: 909–950. DOI: 10.1152/physrev.00026.2013.

33. Tomasi D, Wang GJ and Volkow ND. Energetic cost of brain functional connectivity. Proc Natl Acad Sci U S A 2013; 110: 13642–13647. 20130729. DOI: 10.1073/pnas.1303346110.

34. Bokde AL, Teipel SJ, Drzezga A, et al. Association between cognitive performance and cortical glucose metabolism in patients with mild Alzheimer’s disease. Dement Geriatr Cogn Disord 2005; 20: 352–357. 20050926. DOI: 10.1159/000088558.

35. Xu F, Liu P, Pascual JM, et al. Acute effect of glucose on cerebral blood flow, blood oxygenation, and oxidative metabolism. Hum Brain Mapp 2015; 36: 707–716. 20141016. DOI: 10.1002/hbm.22658.

36. Xie Z, Zhang D, Chung D, et al. Metabolic Regulation of Gene Expression by Histone Lysine beta-Hydroxybutyrylation. Mol Cell 2016; 62: 194–206. DOI: 10.1016/j.molcel.2016.03.036.

37. Kashiwaya Y, Takeshima T, Mori N, et al. D-beta-hydroxybutyrate protects neurons in models of Alzheimer’s and Parkinson’s disease. Proc Natl Acad Sci U S A 2000; 97: 5440–5444. DOI: 10.1073/pnas.97.10.5440.

38. Lim S, Chesser AS, Grima JC, et al. D-beta-hydroxybutyrate is protective in mouse models of Huntington’s disease. PLoS One 2011; 6: e24620. 20110912. DOI: 10.1371/journal.pone.0024620.

39. Wang Z, Li T, Du M, et al. beta-hydroxybutyrate improves cognitive impairment caused by chronic cerebral hypoperfusion via amelioration of neuroinflammation and blood-brain barrier damage. Brain Res Bull 2023; 193: 117–130. 20221226. DOI: 10.1016/j.brainresbull.2022.12.011.

40. Wang N, Yang A, Tian X, et al. Label-free analysis of the beta-hydroxybutyricacid drug on mitochondrial redox states repairment in type 2 diabetic mice by resonance raman scattering. Biomed Pharmacother 2024; 172: 116320. 20240221. DOI: 10.1016/j.biopha.2024.116320.

41. Qiao G, Lv T, Zhang M, et al. beta-hydroxybutyrate (beta-HB) exerts anti-inflammatory and antioxidant effects in lipopolysaccharide (LPS)-stimulated macrophages in Liza haematocheila. Fish Shellfish Immunol 2020; 107: 444–451. 20201105. DOI: 10.1016/j.fsi.2020.11.005.

42. Valdebenito R, Ruminot I, Garrido-Gerter P, et al. Targeting of astrocytic glucose metabolism by beta-hydroxybutyrate. J Cereb Blood Flow Metab 2016; 36: 1813–1822. 20151029. DOI: 10.1177/0271678X15613955.

43. Lehto A, Koch K, Barnstorf-Brandes J, et al. ss-Hydroxybutyrate Improves Mitochondrial Function After Transient Ischemia in the Mouse. Neurochem Res 2022; 47: 3241–3249. 20220608. DOI: 10.1007/s11064-022-03637-6.

44. Zhang L, Dong DL, Gao JH, et al. beta-HB inhibits the apoptosis of high glucose-treated astrocytes via activation of CREB/BDNF axis. Brain Inj 2021; 35: 1201–1209. 20210812. DOI: 10.1080/02699052.2021.1959061.

45. Salman M, Stayton AS, Parveen K, et al. Intranasal Delivery of Mitochondria Attenuates Brain Injury by AMPK and SIRT1/PGC-1alpha Pathways in a Murine Model of Photothrombotic Stroke. Mol Neurobiol 2024; 61: 2822–2838. 20231109. DOI: 10.1007/s12035-023-03739-4.

46. Gureev AP, Sadovnikova IS, Chernyshova EV, et al. Beta-Hydroxybutyrate Mitigates Sensorimotor and Cognitive Impairments in a Photothrombosis-Induced Ischemic Stroke in Mice. Int J Mol Sci 2024; 25 20240524. DOI: 10.3390/ijms25115710.

47. Bazzigaluppi P, Lake EM, Beckett TL, et al. Imaging the Effects of beta-Hydroxybutyrate on Peri-Infarct Neurovascular Function and Metabolism. Stroke 2018; 49: 2173–2181. DOI: 10.1161/STROKEAHA.118.020586.

48. Soto-Mota A, Norwitz NG, Evans RD, et al. Exogenous d-beta-hydroxybutyrate lowers blood glucose in part by decreasing the availability of L-alanine for gluconeogenesis. Endocrinol Diabetes Metab 2022; 5: e00300. 20211116. DOI: 10.1002/edm2.300.

49. Falkenhain K, Daraei A, Forbes SC, et al. Effects of Exogenous Ketone Supplementation on Blood Glucose: A Systematic Review and Meta-analysis. Adv Nutr 2022; 13: 1697–1714. DOI: 10.1093/advances/nmac036.

50. Athinarayanan SJ, Adams RN, Hallberg SJ, et al. Long-Term Effects of a Novel Continuous Remote Care Intervention Including Nutritional Ketosis for the Management of Type 2 Diabetes: A 2-Year Non-randomized Clinical Trial. Front Endocrinol (Lausanne) 2019; 10: 348. 20190605. DOI: 10.3389/fendo.2019.00348.

51. Muller MJ, Paschen U and Seitz HJ. Effect of ketone bodies on glucose production and utilization in the miniature pig. J Clin Invest 1984; 74: 249–261. DOI: 10.1172/JCI111408.

52. Felts PW, Crofford OB and Park CR. Effect of Infused Ketone Bodies on Glucose Utilization in the Dog. J Clin Invest 1964; 43: 638–646. DOI: 10.1172/JCI104949.

53. Ari C, Murdun C, Koutnik AP, et al. Exogenous Ketones Lower Blood Glucose Level in Rested and Exercised Rodent Models. Nutrients 2019; 11 20191001. DOI: 10.3390/nu11102330.

54. Stubbs BJ, Cox PJ, Evans RD, et al. On the Metabolism of Exogenous Ketones in Humans. Front Physiol 2017; 8: 848. 20171030. DOI: 10.3389/fphys.2017.00848.

55. Charoenpiriya A, Chailurkit L and Ongphiphadhanakul B. Comparisons of biochemical parameters and diabetic ketoacidosis severity in adult patients with type 1 and type 2 diabetes. BMC Endocr Disord 2022; 22: 7. 20220106. DOI: 10.1186/s12902-021-00922-3.

