## Supplemental for "β-Hydroxybutyrate enhances brain metabolism in normoglycemia and hyperglycemia, providing cerebroprotection in a mouse stroke model"

**Supplemental File**

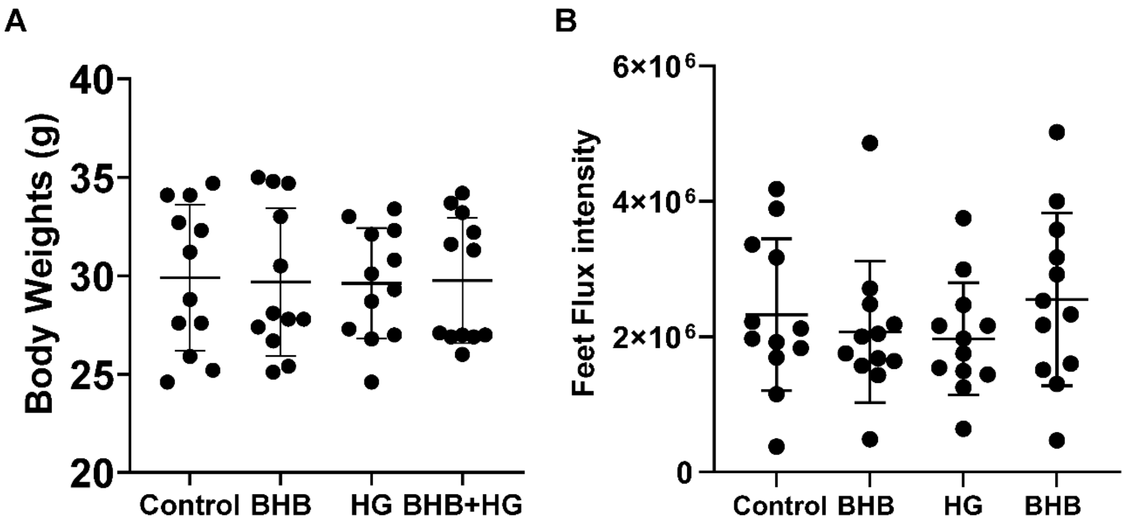


**Supplemental Figure S1. Luciferin-luciferase bioluminescence imaging experiments mice body weights and feet flux intensity.** (**A**) Mice body weights recorded before bioluminescence imaging. (**B**) Total flux emitted from feet in mice 10 minutes post-substrate injection (peak). Data are presented as mean ± SD. Statistical significance was determined using repeated measures one-way ANOVA test followed by Tukey's multiple comparison test showing no significant changes.
